# Similarity index of ex-Gaussian reaction time signatures

**DOI:** 10.1101/2023.05.29.542684

**Authors:** Elias Manjarrez, Angel DeLuna-Castruita, Victoria Lizarraga-Cortes, Amira Flores

**Author notes:** Correspondence: **Corresponding Author: Prof. Elias Manjarrez**, Institute of Physiology, Benemérita Universidad Autónoma de Puebla. 14 sur 6301, Col. San Manuel A.P. 406, C.P. 72570, Puebla, Pue., México, Tels.

## Abstract

In psychology and cognitive neuroscience, reaction time (RT) series and their ex-Gaussian distributions are commonly used to explore the time course of cognitive processes. This study investigated the hypothesis that successive triads of ex-Gaussian mu, sigma, and tau parameters of reaction-time variability across time can be used to construct a geometrical object we termed ex-Gaussian RT-signature, which could help characterize interindividual differences between congruent and incongruent stimuli. To test this hypothesis, we calculated the similarity index of these geometrical objects in young adult participants without detectable neurological disorders. Our findings show that each participant displayed distinct ex-Gaussian RT-signatures in a Cartesian 3D plot, thus exhibiting distinct psychophysical fingerprints. Furthermore, our results revealed that the ex-Gaussian RT-signatures for incongruent stimuli demonstrated a significantly higher similarity index across participants than congruent stimuli (p<0.001; Cohen d=0.4). We suggest that visualizing this psychophysical signature could serve as a valuable tool in characterizing differences in cognitive functioning between individuals, thus providing insights into the development of personalized medicine.

## Introduction

The ex-Gaussian distribution is a powerful tool for modeling reaction time (RT) data in cognitive neuroscience with multiple applications in cognitive testing (Hohle, 1965; Ratcliff and Murdock, 1976; Ratcliff, 1979; Luce, 1986a; Steinhauser and Hubner, 2009; Yamashita et al., 2021; Ging-Jehli et al., 2021). This elegant approach captures the complex and skewed nature of the RT series by convolving a normal distribution with an exponential distribution (Hohle, 1965; Ratcliff, 1979; Heathcote et al., 1991). The ex-Gaussian distribution provides a superior fit compared to the normal distribution. Moreover, the parameters of the ex-Gaussian distribution, mu, sigma, and tau, can disentangle different cognitive processes that contribute to RT performance. The ex-Gaussian distribution paved the way for a deeper understanding of the neural mechanisms underlying cognitive processes. It has been employed many times in the decades since its first application (Hohle, 1965).

However, mu, sigma, and tau parameters could be interpreted with caution (Matzke and Wagenmakers, 2009) in terms of the efficiency, precision, non-decision, and speed of cognitive processes. For instance, Mu represents the central tendency of the reaction time distribution, and it could be related to the rate of the cognitive processes involved in generating a response (Aguilar-Lacasaña et al., 2022). Lower mu values suggest faster and more efficient processing, while higher mu values may represent slower and less efficient processing. Sigma is related to the variability of the reaction time distribution and could be associated with the precision of the cognitive processes involved in generating a response (Chiang et al., 2022). Lower sigma values could indicate more precise and consistent processing, while higher values could indicate less accurate and more variable processing (Galloway-Long et al., 2022). Finally, many authors claim that tau represents the non-decision time of the reaction time distribution and may reflects the time taken for processes that are not directly related to the cognitive processes involved in generating a response (Leth-Steensen et al., 2000; Hervey et al., 2006; Kobor et al., 2015; Kölle et al., 2022). Tau could be related to the speed of these non-decision processes. It may be influenced by factors such as stimulus complexity, motor, and attentional demands, or changes in the efficiency of cognitive processing in that group (Gu et al., 2013).

The abovementioned studies reveal that the different interpretations of the mu, sigma, and tau parameters may have practical uses to examine potential mechanisms related to RT data; however, it would also be helpful to combine these three scalar parameters in a sole geometrical object to define an exclusive feature of each individual. In this context, our study aims to examine whether the conjunction of geometric triads of ex-Gaussian mu, sigma, and tau parameters of reaction-time variability across time represents a distinct “RT signature,” which could be helpful to characterize inter-individual differences. Furthermore, we suggest that by analyzing RT-signatures, researchers could better understand the time course and precision of cognitive processing and the effects of various experimental manipulations on these processes. In this context, our geometrical method and results have significant implications for future research that aims to understand better the neural mechanisms that underlie cognitive functions and could potentially be applied in clinical settings to enhance the diagnosis and treatment of neurological disorders.

## Methods

### Participants

We performed psychophysical experiments on 20 healthy young adults who signed an informed consent form before entering the study. This study was conducted under the principles of the Declaration of Helsinki. A local ethics committee from our university approved the study. A neuropsychologist performed interviews to exclude psychiatric or neurological disorders.

### Stroop test platform and timeline construction

The Stroop test, initially developed by John R. Stroop, was administered to healthy participants as part of the cognitive assessment (Stroop, 1935; MacLeod, 1991). The test involved presenting a colored word to participants, who were instructed to identify the ink color of the word rather than read the word itself. This required inhibition of an automated process (reading the word) instead of a less automatic process (identifying the ink color). We presented the stimuli of the Stroop test in a random sequence of congruent words (words with ink color consistent with the term) or incongruent words (words with a different ink color). In this way, with these random sequences we avoided learning effects. The participants were required to respond to each stimulus as quickly and accurately as possible, with RT recorded for each event. The difficulty of the task varied depending on the coherence of the word, with incongruent stimuli generally resulting in longer RT than congruent stimuli.

We employed the jsPsych framework to develop a Web-based Stroop test which could run online using only a Web browser. The jsPsych library is a powerful JavaScript framework for developing Web-based behavioral experiments (de Leeuw et al., 2023). To develop the software, we first established a timeline for the test, comprising a series of JavaScript trials, each consisting of a plugin designed to perform a specific function. The timeline included the “HTML-keyboard-response” plugin for displaying a welcome message, instructions, and fixation point with a randomized duration (250, 500, 750, 1000, 1250, 1500, 1750, or 2000 milliseconds), as well as the “audio-keyboard-response” plugin for playing a tone at the beginning and end of the test. The “image-keyboard-response” plugin was used to present randomized Stroop test combinations as stimuli after the fixation point, with possible keyboard responses (i.e., ‘y’ for yellow, ‘g’ for green, ‘b’ for blue, and ‘r’ for red) and the corresponding correct or incorrect answer. Time series of RT for correct and incorrect responses were stored in a file.

The software timeline consisted of JavaScript trials and was organized into blocks. The first blocks presented randomly-selected stimuli (congruent and incongruent) with a fixation point before each trial. For example, trial 0 displayed a welcome message, while trials 1-3 displayed the tone and instructions. The final trial provided a summary of the subject’s performance during the test.

### Data Collection and Stimulation Protocol

After the test, the RT data was saved in a .txt or .csv file for further analysis. In each session, we collected 20 congruent and 60 incongruent correct responses per session. To prevent fatigue, the test was divided into three short daily sessions, one in the morning, another in the afternoon, and a third at night. This test-retest protocol was implemented for two active weeks, separated by one week of rest. In this way, for congruent stimuli, we collected 420 RT correct responses during one week and 840 RT correct responses during two weeks; whereas for incongruent stimuli, we got 1260 RT correct responses for one week and 2520 for two weeks. We excluded participants with sleep deprivation because the lack of sleep affects Stroop reaction times (Cain et al., 2011).

### Procedures for dealing with reaction time outliers

Reaction time (RT) outliers are data points that deviate significantly from other observations in a sample. Distinguishing between spurious and genuine RT outliers can be challenging. Spurious outliers are typically influenced by external distractors, lack of interest, sleepiness, fatigue, or unknown events unrelated to the processes under study (Ratcliff, 1993). On the other hand, genuine outliers may be linked to lapses of attention or physiological factors occurring in the brain (e.g., Bender et al., 2015; Adamo et al., 2015; Zhou et al., 2012).

To address this issue, we followed the recommendation proposed by Ratcliff (1993) and examined how the shape of the RT distribution changed with increases in the mean. We employed a test-retest approach, conducting three sessions per day over a seven-day working week, which allowed us to collect small amounts of data per session, but large amounts of data across several days. Subsequently, we aggregated the data across the working week, thereby ensuring a substantial dataset while minimizing fatigue and potential distractors. We repeated this procedure for two weeks separated by a resting week.

We then visualized how the large dataset of RT values changed over time in relation to the mean. Only participants who demonstrated a stable mean value with consistent occurrences of similar RT outliers across the two weeks were included in the final dataset. This approach enabled us to avoid discarding RT outliers that may be meaningful in terms of physiological mechanisms occurring in participants’ brains, thus eliminating concerns associated with setting specific cutoff values for outlier removal.

### Fitting of ex-Gaussian RT distributions

We employed Matlab software (The MathWorks Inc.) (Lacouture and Cousineau, 2008; Johansson, 2023) to fit our RT data for correct responses with ex-Gaussian functions. First, our RT data were plotted as bar histograms to obtain the positively skewed RT distributions. Second, we fitted these RT distributions with the ex-Gaussian function, which is the convolution of the exponential and Gaussian functions. The Matlab code implemented by Lacouture and Coussineau (2008) and Johansson (2023) allows fitting RT distributions in terms of ex-Gaussian probability density functions (PDF) in Matlab. The advantage to use Matlab is that a large community of scientist develops and shares codes making it a powerful programming language. The analytic formula of the ex-Gaussian PDF can be found in many references (Luce, 1986a; Dawson, 1988; Cousineau et al., 2004), for instance in Lacouture and Cousineau (2008); this is the formula:

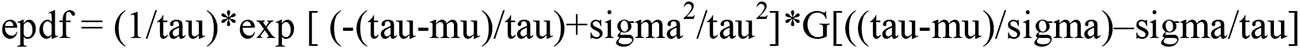

Where epdf is the ex-Gaussian probability density function. A recent Matlab code for this ex-Gaussian epdf formula was developed by Tobias Johansson (2023), which can be found in the exgfit.m file exchange code from MathWorks as follows:

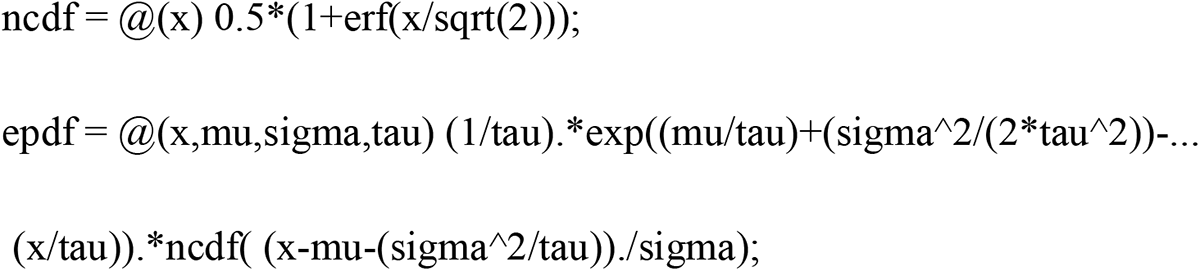

Where [mu,sigma,tau] = exgfit(x,S) fits the ex-Gaussian function to data in vector x using maximum likelihood; whereas ncdf and exp, are the Gaussian and exponential functions, respectively. This code returns the fitted parameters mu, sigma, and tau, where S is a three-element vector of starting values for these mu, sigma, and tau parameters.

The ex-Gaussian PDF (epdf) essentially consists in the exponential function multiplied by the value of the cumulative density of the Gaussian function, which has three parameters mu, sigma and tau. The first two parameters, mu and sigma, correspond to the mean and standard deviation of the Gaussian component, respectively; whereas the third parameter (tau), is the mean of the exponential component.

### Cosine similarity index between ex-Gaussian RT-signatures

We employed the cosine similarity algorithm to calculate the similarity index between two ex-Gaussian RT-signatures (which are two sets of two points in a 3D graph). Note that each ex-Gaussian RT-signature in Figure 2C has two points joined by lines. The cosine similarity measure calculates the cosine of the angle between two vectors, which is a value between -1 and 1. If the angle between the vectors is 0 degrees (i.e., identical), then the cosine similarity is 1. If the angle between the vectors is 90 degrees (i.e., completely different), then the cosine similarity is 0. If the angle between the vectors is 180 degrees (i.e., they are exactly opposite), then the cosine similarity is -1. The following list shows the steps we followed to calculate the cosine similarity between two sets of two points in a 3D graph:

a. First, we calculated the vector between the first two points in each set by subtracting the coordinates of the first point from the coordinates of the second point.
b. Second, we calculated the cosine similarity between the two pairs of vectors using the following formula:

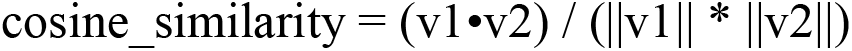

where v1 and v2 are the two vectors being compared, the symbol • represents the dot product of the two vectors, and ||v|| represents the magnitude (length) of the vector. Finally, we calculated the average of the cosine similarities between the two pairs of vectors to get the overall similarity index between the two sets of two points. We normalized the vectors before computing the dot product and magnitude by dividing each vector by its length (i.e., by the square root of the sum of the squares of its components).

Finally, we repeated this procedure for all the combinations of ex-Gaussian RT signatures in our study. Hence, we obtained a similarity index matrix and the corresponding color map in the range [-1, 1] to show the full range of the similarity index from -1 to 1 for all the possible combinations between pairs of participants.

### Statistical analysis

For each control participant, we analyzed the correct responses to congruent and incongruent stimuli of the Stroop task over two weeks. We analyzed for each week 420 RT values of correct responses for congruent and incongruent stimuli for each individual. We selected the same number of 420 RT values of correct responses for congruent and incongruent stimuli in order to make fair comparisons between both conditions and avoid bias due to different sample size. We also chose this number of RT responses, given that at least 100 RT observations per subject are required to obtain reasonable ex-Gaussian parameter estimates (Ratcliff, 1979), which are affected with a low sample size below 100 (Lacouture and Cousineau, 2008). We got histograms of these RT series, which were fitted with ex-Gaussian functions using Matlab. Hence, we obtained the mu, sigma, and tau ex-Gaussian weekly parameters for each participant in the congruent and incongruent conditions. With these groups of mu, sigma, and tau ex-Gaussian parameters, we constructed geometrical objects termed ex-Gaussian RT-signatures and calculated the cosine similarity index distributions (see results section).

We compared the similarity index distributions of ex-Gaussian RT-signatures for congruent and incongruent stimuli. Then we performed the non-parametric Mann-Whitney U test following a Kolmogorov–Smirnov normality test, p<0.05. We examined statistically significant differences between the similarity-index distributions of RT-signatures between ex-Gaussian RT-signatures for congruent and incongruent stimuli. Moreover, we computed the effect sizes for group comparisons using Cohen’s d (Cohen, 2013). Finally, we performed the statistical analysis with SPSS Sigma Plot software.

## Results

In the upper panel of Figure 1A, we show the raw RT data obtained from Participant #7 during the presentation of congruent stimuli of the Stroop task. These sequences of RT values were obtained over two weeks, as illustrated. The lower panel of Figure 1A illustrates the corresponding RT distributions in color gray and the fitted ex-Gaussian probability density functions in color magenta. The insets in this lower panel show the best-fitting values of sigma, tau, and mu ex-Gaussian parameters, which change over the two weeks. In the same way, we display in the upper panel of Figure 1B RT data obtained from Participant #7, but during the presentation of incongruent stimuli of the Stroop task. Note that RT values and ex-Gaussian parameters for the incongruent stimuli exhibit more significant variability than those for the congruent stimuli depicted in panel A.

**Figure 1.**
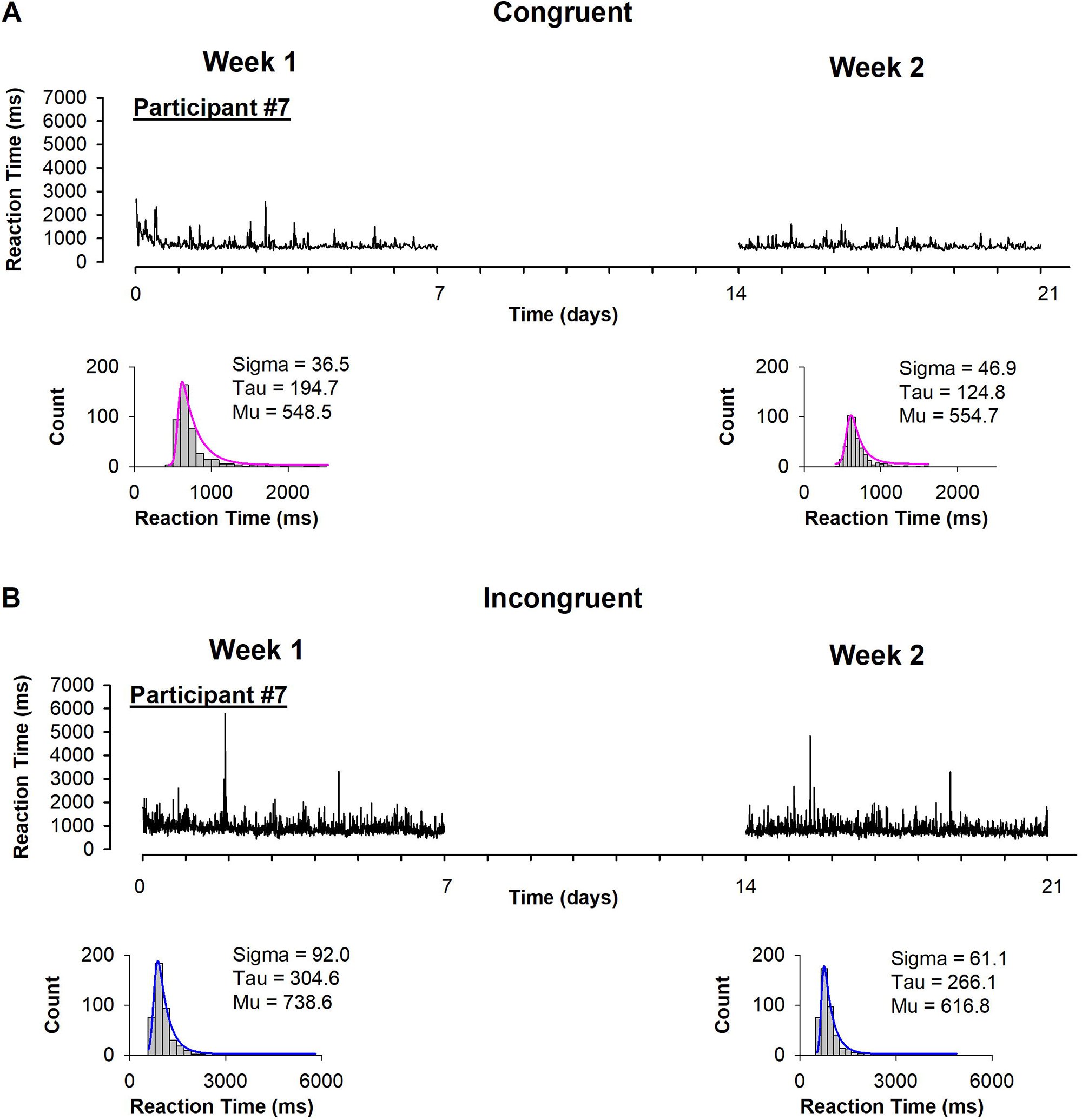
Example of RT raw data, distributions, and ex-Gaussian best fitting of probability density functions (PDF) for congruent (**A**) and incongruent (**B**) stimuli for participant #7.

We employed the triads of ex-Gaussian RT parameters: x=sigma, y=tau, and z=mu, obtained during two weeks in Figure 1A and Figure 1B to construct the 3D geometrical objects in coordinates (x,y,z) illustrated in Figure 2A and Figure 2B, respectively. Because these geometrical objects were obtained from the ex-Gaussian parameters and they resemble “signatures”, we termed them “ex-Gaussian RT-signatures.” If we compare Figure 2A and Figure 2B, we can note that the ex-Gaussian RT-signatures associated with the congruent and incongruent stimuli for this participant #7 are different.

**Figure 2.**
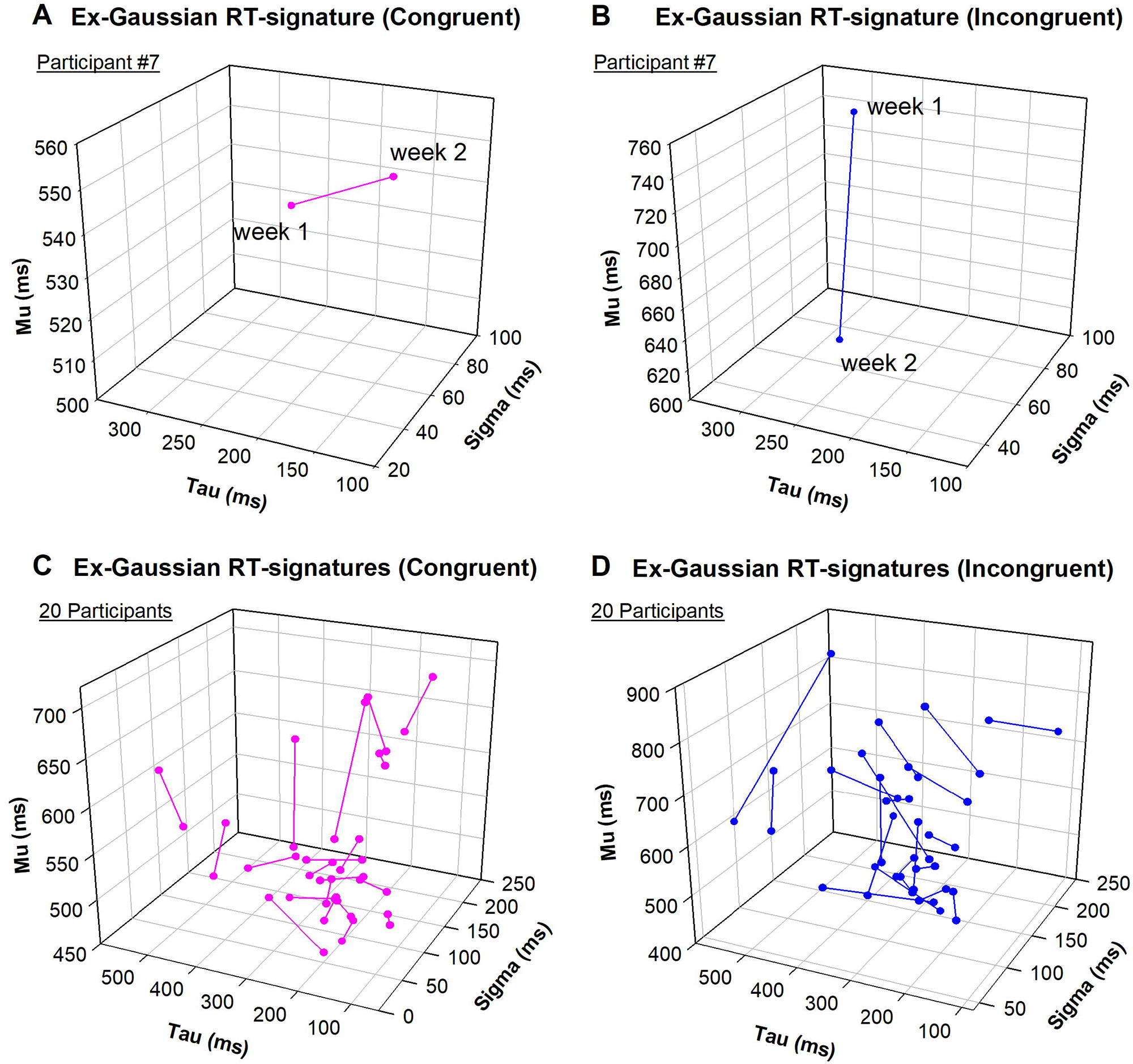
Ex-Gaussian RT signatures for congruent and incongruent stimuli. **A and B**, ex-Gaussian RT signatures for congruent and incongruent stimuli for participant #7, respectively. **C and D**, the same as A and B but for 20 participants.

Hence, we applied the same procedure described in the previous two paragraphs, illustrated in Figures 1A, 1B, 2A, and 2B, to all the participants in this study. The results are visualized in Figures 2C and 2D, which show pooled data of these ex-Gaussian RT signatures. An inspection of Figures 2C and 2D qualitatively highlights interindividual differences in the ex-Gaussian RT-signatures among all the participants for congruent and incongruent stimuli. We also can observe that the ex-Gaussian RT signatures for congruent stimuli (Figure 2C) exhibit different configurations in the 3D space than those for congruent stimuli (Figure 2D).

Finally, we performed a quantitative analysis to compare the similarity among the ex-Gaussian RT-signatures illustrated in Figures 2C and 2D. First, we calculated the cosine similarity index among the ex-Gaussian RT-signatures for congruent stimuli obtained for all the participants; for instance, we compared the similarity index (see Methods section) of ex-Gaussian RT-signatures between participants #1 and #2, #1 and #3, and so on, to complete all the possible combinations among all pairs of subjects in Figure 2C. In Figure 3A and Figure 3C, we show the quantitative results of these similarity indexes of ex-Gaussian RT-signatures for congruent stimuli. Second, we calculated the cosine similarity index among the 3D ex-Gaussian RT-signatures for incongruent stimuli obtained for all the participants in Figure 2D. In Figure 3B and Figure 3D, we show the quantitative results of these similarity indexes of ex-Gaussian RT-signatures for incongruent stimuli.

**Figure 3.**
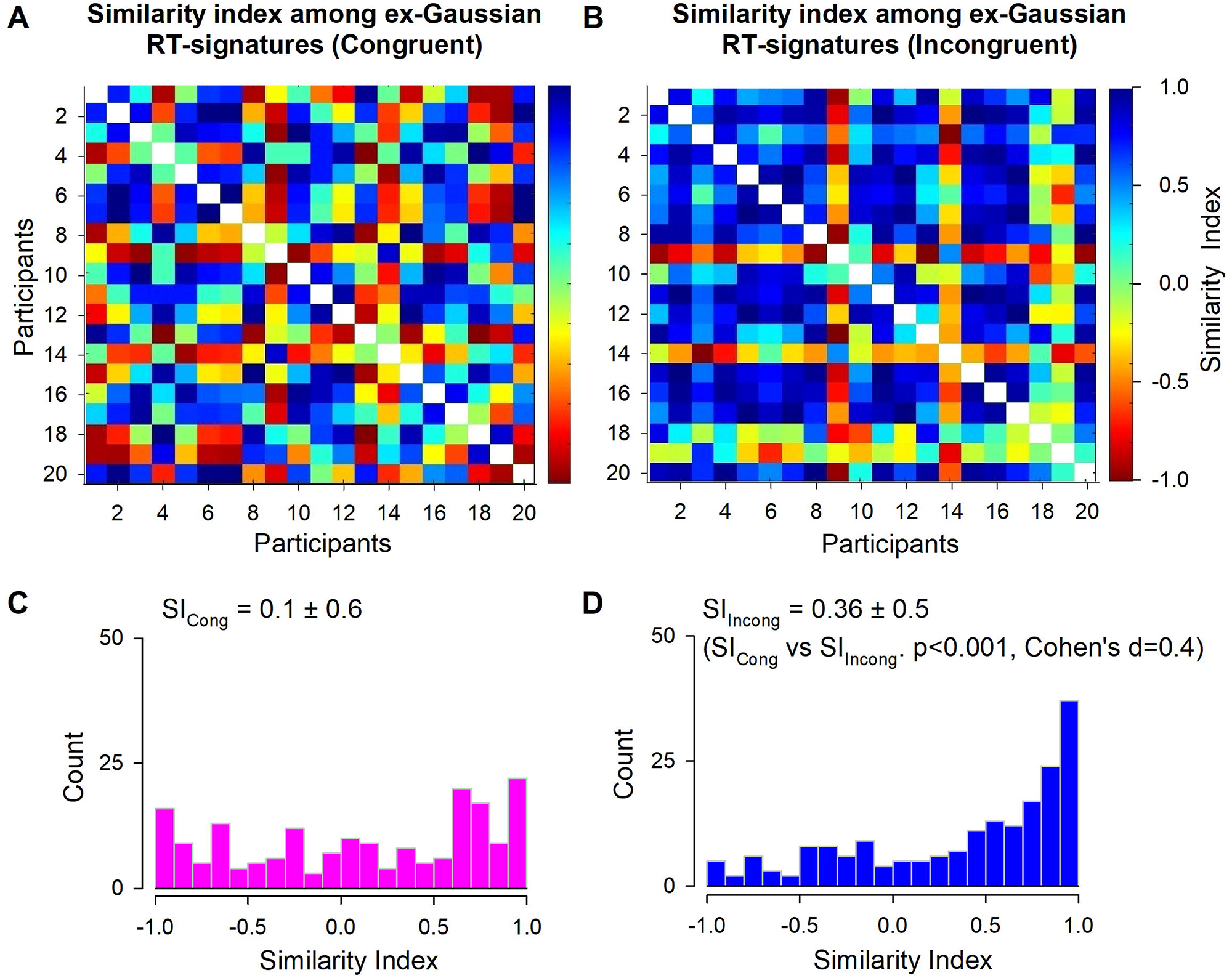
Results of the similarity index analysis of ex-Gaussian RT signatures for all the participants. **A and B**, similarity index maps for the ex-Gaussian RT signatures. **C and D**, histogram distributions for the similarity index of ex-Gaussian RT signatures for all the participants for congruent and incongruent stimuli, respectively.

A qualitative inspection of Figure 3 reveals a more significant number of participants with a greater similarity index of ex-Gaussian RT-signatures for incongruent stimuli (see the number of blue squares in Figure 3B) than congruent stimuli (see the number of blue squares in Figure 3A). If the cosine similarity measure (see Methods section) is one, then the ex-Gaussian RT-signatures are identical (blue squares). Conversely, if the cosine similarity is 0, then the ex-Gaussian RT-signatures are entirely different (green squares); in the same way, if the cosine similarity is -1, then the ex-Gaussian RT-signatures also are entirely different and are exactly opposite (red squares). In this context, note that Figure 3 reveals a more significant number of participants with a lower similarity index of ex-Gaussian RT-signatures for congruent stimuli (see the high number of green and red squares in Figure 3A) than for incongruent stimuli (see the low number of green and red squares in Figure 3B).

A quantitative analysis of Figure 3C and 3D reveals that the mean similarity index (SI) for the ex-Gaussian RT signatures of incongruent stimuli across all participants is 0.36 ± 0.5. In contrast, the mean similarity index for the ex-Gaussian RT signatures of congruent stimuli across all participants is 0.01 ± 0.6. A statistical comparison demonstrates that the mean similarity index for incongruent stimuli is significantly different from that of congruent stimuli (p<0.001, Mann-Whitney U test, N=20 participants, n=190 similarity indexes, Cohen’s d=0.4).

## Discussion

We introduced for the first time the visualization of a psychophysical metrics termed “ex-Gaussian RT-signature”, which is composed by the sigma, tau, and mu parameters of the ex-Gaussian distribution measured at different consecutive times. Our findings show that participants displayed distinct ex-Gaussian RT-signatures in a Cartesian 3D plot; i.e., exhibiting distinct “psychophysical fingerprints”. Furthermore, we found that the similarity indexes of these ex-Gaussian RT-signatures among participants are significantly greater for incongruent stimuli than congruent stimuli. This highlights that healthy participants exhibit more significant interindividual differences in their normal cognitive functioning when processing congruent stimuli than incongruent stimuli.

We employed the similarity-index metrics to compare the ex-Gaussian RT signatures because it is a way to measure how similar two things are to each other. The similarity index algorithms have gained popularity recently thanks to the advances in the artificial intelligence (AI) field (Brown et al., 2020), which could have important applications in a generalist medical AI (Moor et al., 2023). Similarity index algorithms are used in fields such as data science, machine learning, complex network analysis, and information retrieval to compare stuff like text documents or images (Chang et al., 2012; Huebner and Gegenfurtner, 2012; Spano et al., 2020; Lu et al., 2009; Wang et al., 2016). In psychology and neuroscience, the similarity index is also a helpful tool (Lu et al., 2009; Gerstorf et al., 2013; Vinding et al., 2016; Bouts et al., 2017; Doucet et al., 2020; Urriola et al., 2020; Pitkänen et al., 2020; Meijer et al., 2020; Tian et al., 2021; Janssen et al., 2021; West et al., 2022; Baldwin et al., 2022; Xuan-Tran et al., 2022; Sari et al., 2023; Barca et al., 2023).

In general, a distance metric is one common way to measure similarity (Lu et al., 2009). A distance metric measures how far apart two things are from each other, with a smaller distance indicating a higher level of similarity. The cosine similarity measure is one example of a distance metric often employed in medical and biological fields (Wang et al., 2016; Voorneveld et al., 2018; Nath et al., 2021; Mofidi et al., 2022). Due to these previous applications, we used cosine similarity to calculate the similarity index between two sets of two points (the ex-Gaussian RT-signatures for a pair of participants) in a 3D graph; however, our analysis could be extended to a greater number of points (more weeks), as explained in the next paragraph.

In this study, our ex-Gaussian reaction time (RT) signature is represented by two distinct points, each corresponding to a specific week, with coordinates (sigma, tau, mu). The observation that these points differ for each participant indicates the presence of intraindividual variability across the two-weeks period. However, it is possible to construct ex-Gaussian RT signatures with additional points, encompassing a greater number of weeks. Furthermore, it is also feasible to explore alternative time scales for each point, such as using a day-scale or a month-scale instead of the week-scale employed in this study. However, it is essential to exercise caution when adopting different time scales, as the RT sample size should exceed n=100 to ensure the validity of the ex-Gaussian parameters (Ratcliff, 1979; Lacouture and Cousineau, 2008). In our case we employed a RT sample size of 420 for every point of the ex-Gaussian signature to ensure the validity of the ex-Gaussian parameters.

The intraindividual variability across time revealed by the ex-Gaussian RT signatures constructed across two weeks confirms previous studies that intraindividual variability is an intrinsic phenomenon occurring in healthy subjects. In fact, Luce (1986) expressed his concern that we cannot estimate just one RT distribution because in fact there are many RT distributions, and what we report as an estimate of a distribution really is an estimate of a probability mixture of many distributions. In this context, the ex-Gaussian RT signatures taken at different time scales, as mentioned in the previous paragraph, can explain the concern made by Luce, and therefore they may be helpful to accurately characterize the evolution of RT distributions across time per individual. A perspective of our present research is that future studies could examine ex-Gaussian RT signatures at different time scales and their relationship with different cognitive processes.

Additionally, the utilization of ex-Gaussian RT signatures presents an opportunity to investigate sequential effects in “memory” search designs of RT distributions. This could be relevant given that the sequential effects could also represent a concern as outlined in the caution words by Luce (1986). For instance, by segregating the RT data based on the stimulus-response histories (i.e., “memory” sets) it could be possible to analyze and visualize the inherent sequential effects by means of the ex-Gaussian RT signatures. Hence, by incorporating ex-Gaussian RT signatures, researchers could effectively address and account for these inevitable sequential effects instead of disregarding them.

The ex-Gaussian RT signatures could serve as a valuable tool for examining functional mechanisms associated with the RT intraindividual variability in both healthy individuals and patients with neurological disorders. In this context, it would be interesting to re-examine previous studies describing a relationship between the intraindividual RT variation and the variability of single trial brain activation (Bender et al., 2015; Adamo et al., 2015; Zhou et al., 2012), but in the context of the ex-Gaussian RT signatures. This could be relevant, for instance, to explore why the findings from trial-by-trial analyses of RT intraindividual variability over time (Bender et al., 2015; Adamo et al., 2015; Zhou et al., 2012) contradict other analyses conducted on average procedures of mean RT, which claimed moderate to high levels of reliability for RT measurements over time (Liu et al., 2017).

The significance of our study is that the ex-Gaussian RT signature could be beneficial in different fields of cognitive neuroscience that use ex-Gaussian distributions to examine intra or interindividual variability in RT data. To support this claiming, there are reports of previous applications of the ex-Gaussian analysis in various populations, including patients with schizophrenia (Fish et al., 2018; Panagiotaropoulou et al., 2019), bipolar disorder (Gallagher et al., 2015; Vainieri et al., 2020; Burgess et al., 2022), lexicality effects in word recognition (Balota et al., 1999), and cocaine-dependent subjects (Liu et al., 2012). Additionally, researchers have utilized the ex-Gaussian distribution to investigate the interplay between state anxiety, heart rate variability, cognition (Spangler et al., 2018, Fitousi et al., 2020; 2021; 2022), and ADHD (Kofler et al., 2013; Ging-Jehli et al., 2021).

In conclusion, the ex-Gaussian distribution and its parameters visualized in the form of a psychophysical signature could provide a powerful tool for investigating the underlying cognitive processes of RT performance and also could have the potential to inform the development of personalized interventions for individuals with cognitive impairments. Furthermore, this psychophysical ex-Gaussian RT signature may represent a neuronal feature of every participant (i.e., a psychophysical fingerprint) that could be utilized in future studies to develop novel metrics to identify individuals.

## Acknowledgments and funding

The following grants supported this research: Cátedra Marcos Moshinsky (EM), CONAHCYT-SNI (EM), Dirección de Internacionalización de la Investigación (EM), and Proyectos-VIEP-PRODEP-Cuerpos-Académicos-BUAP-Puebla (EM), México.

